# Low-frequency neural responses synchronize to distinct structural rather than lexical features during sentence comprehension

**DOI:** 10.64898/2026.08.10.743974

**Authors:** J. Martorell, S. Mancini, P.M. Paz-Alonso, M. Carreiras, N. Molinaro

## Abstract

Language comprehension involves the integration of single words (lexical units) into phrases and sentences (multi-word structures). Previous frequency-tagging studies have found that low-frequency neural responses synchronize to the frequency of multi-word structures. However, it is currently unclear how exactly structural and lexical processes jointly impact these synchronization findings. The present magnetoencephalography experiment implemented the frequency-tagging paradigm in the visual modality with written words to investigate neural synchronization to multi-word sentences varying in internal structure (reversed word orders between verb-initial Spanish and verb-final Basque sentences) and in lexical content (real words and pseudo words). We find converging evidence that neural responses largely synchronize to structural rather than lexical features. This was observed as robust phase synchronization strength to the frequency of sentences containing reversed structures, with certain lexical modulations depending on language-specific structural features. Crucially, we also found shifted phase angle dynamics between the reversed structures of Spanish and Basque sentences independently of word-level lexical characteristics. Together, these findings suggest that neural synchronization to multi-word structures is largely driven by distinct structural features operating via two segregated neural dimensions: frequency coding for the coarser aspects (i.e., timescale/duration) and phase representing the finer-grained aspects (i.e., internal structure) of multi-word structures. Our findings thus advance key insights into the core components of the neural mechanisms supporting language comprehension.

**Highlights:**

- Neural synchronization to sentences is driven by structural (not lexical) features.
- Robust sentence-frequency synchronization across languages varying in structure.
- Phase angle is selectively sensitive to cross-linguistic structural differences.
- Lexical modulations depend on language-specific structure.
- Structure synchronization segregates into two dimensions: frequency and phase.

## Introduction

Language comprehension involves complex cognitive processes mapping bottom-up sensory cues (e.g., speech acoustics or written words) to top-down abstract representations (Bever and Poeppel, 2010; Martin, 2016, 2020; Meyer et al., 2020a, 2020b). Such sensory-to-abstract mapping processes support the segmentation of continuous sensory input into single words representing specific lexical meanings. Together with single-word lexical meaning, language comprehension is additionally achieved by integrating single words into coherent multi-word abstract structures such as phrases and sentences. Electro- and magnetoencephalography (EEG & MEG) evidence suggests that such multi-word structural processes rely on neural mechanisms exhibiting selective spatio-temporal dynamics at varying timescales across fronto-temporal regions (see (Friederici, 2017; Meyer, 2018; Hagoort, 2019; Pylkkänen, 2019, 2020; Martorell et al., 2023)). Neural responses at low frequencies (< 4 Hz) have notably been linked to structural processes using the frequency-tagging paradigm (Buiatti et al., 2009; Nozaradan et al., 2011; Nozaradan, 2014). Specifically, it has been shown that neural responses synchronize to the frequency (i.e., timescales) of abstract multi-word structures lacking sensory correlates (e.g., pitch or intonation cues) in the stimulus ((Ding et al., 2016, 2017b; Sheng et al., 2018; Meng et al., 2020); for reviews, see (Meyer, 2018; Zoefel et al., 2018; Martorell et al., 2023)). Such frequency-tagging findings are generally linked to top-down abstract responses beyond the bottom-up sensory processing of the stimulus (Meyer et al., 2020a, 2020b; Ding, 2025; Martorell et al., 2026). However, there is ongoing debate regarding the precise roles of structural and lexical processes in generating these abstract-level neural responses.

Low-frequency neural responses synchronized to multi-word structures have intuitively been interpreted as largely reflecting structural processing (Ding et al., 2016, 2017b; Ding, 2023a, 2025). The dominant role of structural processing has been challenged by computational findings showing that single-word lexical information is sufficient to elicit such synchronized responses ((Frank and Yang, 2018); cf. (Martin and Doumas, 2017)). In line with this lexical explanation, sentences with identical lexical content but subtle structural differences (e.g., genitive vs. dative morphosyntax) result in indistinguishable neural responses (Kalenkovich et al., 2022). Yet, lexical processing cannot apparently account for other neural evidence observed in distinct structural contexts that keep constant word-level information. For example, neural responses to sentences are disrupted when the same words appear in reversed orders representing incorrect structures (Lo et al., 2022). Likewise, neural responses are sensitive to sentences containing identical words in distinct word orders (subject-verb-object vs. object-subject-verb) representing correct structures (Chen et al., 2025). These findings are instead consistent with a modulatory rather than driving role of single-word lexical processing as also suggested by other frequency-tagging studies (Burroughs et al., 2021; Zhao et al., 2024; Xie et al., 2025). Together, current evidence is mixed about how structural and lexical components precisely contribute to the neural synchronization to multi-word structures (for discussion, see (Ding et al., 2017a; Frank and Christiansen, 2018; Martorell et al., 2023)).

Isolating the contribution of structural components is challenging as structural manipulations often vary with other non-structural aspects (see (Pylkkänen, 2019, 2020)). This is likely the case even for the reported word-order contrasts (Lo et al., 2022; Chen et al., 2025) despite yielding neural patterns difficult to reconcile with a purely lexical explanation. Incorrect structures (as in (Lo et al., 2022)) introduce error detection and monitoring processes, while within-language word-order comparisons (as in (Chen et al., 2025)) vary statistical distribution patterns (e.g., certain word orders are more frequent than others) and convey distinct pragmatic meanings at the discourse level (e.g., focused information). Importantly, this sort of non-structural processes often elicits neural responses exhibiting slow dynamics (i.e., P600 component, see (Schlesewsky et al., 2003; Bornkessel-Schlesewsky and Schlesewsky, 2009; Erdocia et al., 2009; Molinaro et al., 2011; Regel et al., 2014; Dröge et al., 2016; Yano and Koizumi, 2018)), which can interact with structural processes linked to low-frequency neural responses ((Meyer, 2018; Henke and Meyer, 2021); for further discussion, see (Martorell et al., 2023)). Such non-structural confounds can be avoided by instead comparing typologically distinct languages that naturally differ in their default word order. For example, a widespread cross-linguistic typological difference is the relative position of verbs and their arguments (e.g., direct-object nouns), which leads to the separation between verb-initial (verbs precede their arguments, e.g., English and Spanish) and verb-final (arguments precede their verbs, e.g., Japanese and Basque) languages (see (Kayne, 1994; Newmeyer, 2004; Hawkins, 2004; Abels and Neeleman, 2012; Haider, 2015; Levshina, 2019; Levshina et al., 2023)). Unlike other word-order manipulations, comparing internally reversed structures between verb-initial and verb-final languages involves a structural manipulation without additional non-structural confounds (i.e., no errors involved and similar statistical distributions and pragmatic meanings between languages). Furthermore, in line with other language-specific incremental processing dynamics depending on word order (e.g., (Rubio-Fernandez and Jara-Ettinger, 2020; Morucci et al., 2024)), verb-final languages seem to rely more strongly on predictive processes (Levy and Keller, 2013; Husain et al., 2014; Coopmans et al., 2025) as initial words containing informative lexical cues (e.g., content words representing arguments) can be used earlier to anticipate both lexical and structural features (e.g., specific verbs and morphosyntactic cues from verbs, respectively) relevant for comprehension. This might also result in stronger lexical modulations for verb-final than verb-initial languages, thereby reflecting how the relative sensitivity to lexical processing depends on structural considerations. Such cross-linguistic approaches can thus provide a more direct estimation of how structural and lexical processes jointly contribute to multi-word neural synchronization. This would also broaden the empirical coverage of frequency-tagging findings, with important implications for the fundamental characterization of language comprehension mechanisms from a cross-linguistic perspective (see (Bornkessel-Schlesewsky and Schlesewsky, 2013, 2016; Norcliffe et al., 2015; Coopmans et al., 2025)).

The contribution of lexical processing is often assessed by manipulating word-level information. A productive word-level approach from single-phrase or single-sentence paradigms is the use of the so-called Jabberwocky structures in which content words are replaced by pseudo words that resemble real words in that language (Pallier et al., 2011; Fedorenko et al., 2016; Kaufeld et al., 2020; Coopmans et al., 2022; Rafferty et al., 2023, 2024; Shain et al., 2024). Because abstract structure can be inferred from relevant morpho-syntactic cues (e.g., function words and subject-verb agreement and case morphology), this sort of word-level approach allows for evaluating whether neural responses are more sensitive to structural information (present in both real- and pseudo-word structures) or lexical information (preserved in real-word but considerably reduced in pseudo-word structures). For example, it has been shown that the presence of lexical content can result in enhanced time-resolved neural synchronization to abstract structure ((Kaufeld et al., 2020; Coopmans et al., 2022); cf. (Rafferty et al., 2024)). Besides a computational study (Martin and Doumas, 2017), it is noteworthy that frequency-tagging experiments have not yet used this sort of word-level approach to disentangle how structural and lexical components impact the neural synchronization to multi-word structures.

Together with linguistic manipulations, the examination of temporal dynamics can provide crucial evidence about the neural synchronization to structural and/or lexical components (Nelson et al., 2017; Kaufeld et al., 2020; Coopmans et al., 2022; Slaats et al., 2024; Weissbart and Martin, 2024; Coopmans et al., 2025). This is because neural responses aligned to each of these components should follow their distinct timescales: present across the multiple words of the structure and/or most prominent at specific single words. Although it has been reported that low-frequency neural activity follows the timescales of certain multi-word structures rather than single words (Ding et al., 2016; Zhang and Ding, 2017), the vast majority of frequency-tagging studies have only focused on conventional frequency-resolved analysis (e.g., power or inter-trial phase coherence spectra) that disregard the temporal dimension. Such conventional frequency-domain analyses assess phase synchronization strength and can effectively dissociate neural dynamics occurring at different frequencies but, crucially, have well-known limitations related to their poor sensitivity to temporal dynamics (see (Zhou et al., 2016; Ding, 2023b; Zhang et al., 2023)). For example, two neural signals occurring at the same frequency with comparable magnitude (i.e., amplitude) but temporally reversed dynamics would result in identical outcomes. This means that conventional frequency-domain analyses would fail to capture the reversed dynamics of neural responses following the internal structure of reversed sentences (e.g., verb-initial vs. verb-final structures). Interestingly, it has been shown that the phase angle of low-frequency neural responses dynamically varies depending on how listeners decide to segment structurally ambiguous sentences into shorter/longer phrases ((Meyer et al., 2016); see also (Henke and Meyer, 2021)). This and other links between time-resolved phase shifts and top-down modulations (Lakatos et al., 2008; Besle et al., 2011; Ten Oever and Sack, 2015; Ten Oever et al., 2024) clearly indicate the relevance of phase angle to elucidate how neural dynamics support the inference of abstract information during language comprehension. Indeed, by additionally considering phase angle, it could be possible to segregate the simultaneous processing of distinct linguistic features into different neural dimensions (e.g., frequency and phase; see also (Martorell et al., 2023)). This sort of multi-dimensional approach could thus reveal novel insights into how neural responses synchronize to structural and lexical features.

In this MEG study, we investigated how structural and lexical processes simultaneously contribute to the neural synchronization to multi-word sentences. Specifically, abstract structure was manipulated from a cross-linguistic perspective (comparing internally reversed word orders between verb-initial sentences in Spanish and verb-final sentences in Basque) and lexical content by contrasting sentences containing either real or pseudo words (i.e., Jabberwocky sentences). To rule out additional confounds related to the between-language comparison, the experiment was implemented in the same group of Spanish-Basque balanced bilinguals with high proficiency in both languages. Using the frequency-tagging paradigm in the visual modality (see (Pei et al., 2023)), we presented written words that corresponded to multi-word sentences occurring at identical frequencies in two internal structures (verb-initial in Spanish or verb-final in Basque) and two word types (real words or pseudo words). We assessed neural synchronization to multi-word sentences using two distinct analytical approaches: (i) Inter-Trial Phase Coherence (ITPC) to capture a more generic frequency-domain measure of phase synchronization strength and (ii) phase angle to obtain a more precise estimation of temporal dynamics. Given the mentioned limitations of frequency-domain approaches, we expected that ITPC might not necessarily result in distinct phase synchronization patterns for the reversed structures from Spanish and Basque sentences presented at identical frequencies. Based on prior evidence (Meyer et al., 2016), we hypothesized that phase angle could instead capture this neural sensitivity to structural features, specifically by reflecting shifted phase angle dynamics (potentially approximating anti-phase relationships) between the internally reversed structures of Spanish and Basque sentences. Moreover, apart from testing whether lower lexical content generally decreases neural synchronization to sentences (i.e., real words > pseudo words) similarly to findings from other paradigms (Kaufeld et al., 2020; Coopmans et al., 2022), the simultaneous manipulation of structure and word type would allow us to clarify the relative contribution of structural and lexical processes. More concretely, given their verb-final structural features, Basque sentences might be more sensitive to lexical information than Spanish sentences, possibly reflected as stronger phase synchronization for real than pseudo words only in Basque. Taken together, our MEG study aimed to provide a more exhaustive characterization of the core components underlying the neural mechanisms of language comprehension.

## Methods

### Participants

26 Spanish-Basque balanced bilingual participants participated in the experiment. Data from 2 participants could not be recorded due to technical problems with the MEG system and data from 2 other participants were discarded due to excessively noisy recordings. This resulted in a final sample of 22 participants (16 female; mean age = 29.27 years, standard deviation = 6.84; range: 21-43). Sample size could not be increased due to resource constraints (Lakens, 2022), particularly considering the requirement of participants with specific linguistic profiles (see below). Participants were right-handed and reported no history of neurological, hearing, or language disorders. They had normal or corrected-to-normal vision. The experiment was approved by the Basque Center on Cognition, Brain and Language (BCBL) ethics committee. Written consent was obtained from participants before participation.

Participants were highly proficient in both Spanish and Basque, although they showed certain variability in their bilingual profile. First, their language proficiency was estimated by the BEST test (de Bruin et al., 2017), which provides scores (maximum: 65) from a picture-naming task for each language. Participants’ BEST scores were slightly more variable in Basque (mean: 62.09; SD: 2.89; range: 56-65) than in Spanish (mean: 64.59; SD: 0.96; range: 61-65). Second, based on self-reported language dominance, Spanish was the dominant language in 14 participants and Basque in 8 participants. Third, considering their relative Age of Acquisition (AoA) between languages, 6 participants first acquired Spanish, 5 Basque, and 11 both Spanish and Basque from birth.

### Materials

Experimental materials comprised multi-word sentences with reversed word orders between Spanish and Basque sentences (see Figure 1A). More concretely, sentences were composed of 3 visual chunks containing written words (each from a different word class): an auxiliary verb, a verb (in past participle form), and a Noun Phrase (NP) in Spanish (e.g., [*hemos*] [*vendido*] [*una casa*]); an NP, a verb (in past participle form), and an auxiliary verb in Basque (e.g., [*etxea*] [*saldu*] [*dugu*]). Stimuli were chosen arbitrarily for each language such that they were not translations between Spanish and Basque (shown in Figure 1 only for visualization purposes; see Table S1 for descriptive statistics of psycholinguistic variables). Words were always different across sentences except for the auxiliary verb, which was always the same word (in Spanish: *hemos*; in Basque: *dugu*). In addition to these sentences with real words, the experiment included multi-word sentences containing pseudo words in both Spanish and Basque (see Figure 1A). Pseudo-word sentences were created by substituting content words (nouns and verbs) and keeping function words (auxiliary verbs in Spanish/Basque; article determiner in Spanish NPs) along with morphosyntactic information (i.e., morphological suffixes to nouns and verbs). Specifically, pseudo words for nouns and verbs were created from the Spanish/Basque stimuli selected for the real-word conditions using the software Wuggy (Keuleers and Brysbaert, 2010). Pseudo words were of the same length (in characters) as their corresponding real-word conditions in Spanish/Basque (see Table S1). This resulted in pseudo-word sentences with lower lexical content but similar structural features as the real-word conditions, also respecting the Spanish/Basque phonotactic rules.

**Figure 1.**
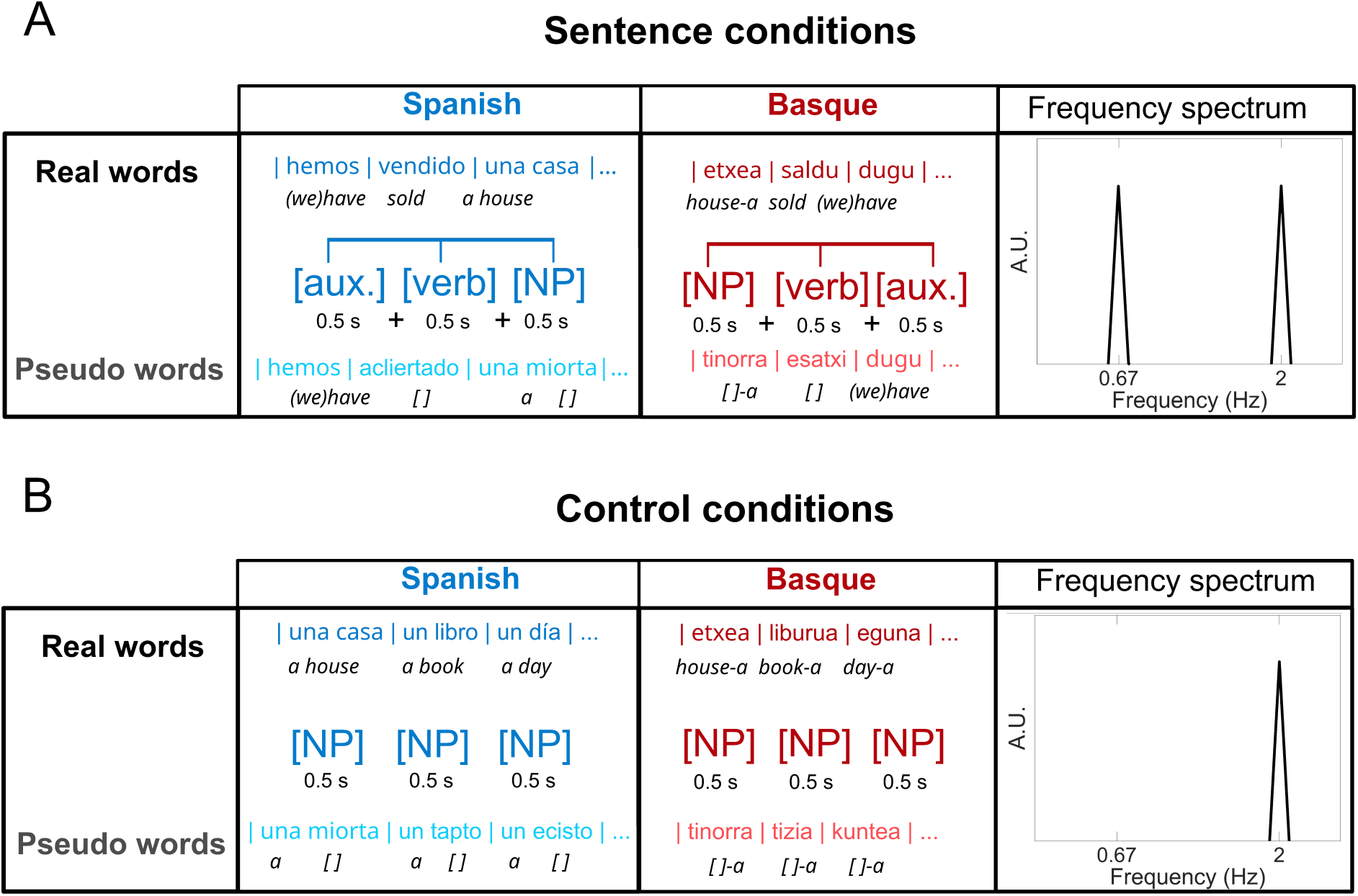
Experimental design of frequency-tagging study in the visual modality (presenting written words). **A)** Sentence conditions corresponding to 3-word sentences (containing an auxiliary verb, aux; a (main) verb; and a Noun Phrase, NP) in two languages (Spanish in blue; Basque in red) and in two word types (Real words: top row in darker colors; Pseudo words: bottom row in lighter colors). Middle row represents the sentence structure, with each word inside a bracket corresponding to a written word presented as a single visual chunk. Note the reversed word order between Spanish and Basque sentences. Rightmost panel includes the frequency spectrum (X-axis = frequency in Hertz, Hz; Y-axis = representative peak value in arbitrary units, A.U) representing the frequencies of interest shared across conditions given the duration of single words (0.5 s = 2 Hz) and 3-word sentences (0.5 s x 3 = 0.67 Hz). **B)** Control conditions corresponding to single words (NPs) without abstract structure between them in the same languages and word types as the sentence conditions (see A). Note that the frequency spectrum only displays a peak at the single-word frequency (2 Hz), unlike the sentence conditions (see A).

Unlike the words corresponding to the verb and NP, the auxiliary verb word was always identical in Spanish and Basque sentences (i.e., *hemos* and *dugu*, respectively). Note that the Spanish NPs contained two words representing an article and a noun (e.g., *una casa*), while Basque NPs contained one word with the article attached to the noun as a suffix (e.g., -*a* in *etxea*), thereby resulting in different lengths in terms of number of words as well as characters (see Table S1). This length difference for NPs is intrinsic to the properties of each language in their written form and could not be controlled given that the experiment was implemented in the visual modality. To control for this salient perceptual (visual) difference in length between Spanish and Basque NPs, we additionally included a control condition containing similar NPs in each language and lexical type (see Figure 1B). This control condition only included NPs that did not require further syntactic structuring with other NPs. The experiment included additional conditions consisting of 2-word verb phrases, which are not reported in the present study as they address different theoretical questions.

### Procedure

Participants were visually presented with the sequences of written words projected on a screen. Stimuli were presented via a PROPixx projector (VPixx Technologies Inc.) using Psychtoolbox (Kleiner et al., 2007) in Matlab. These sequences were presented in trials (12 per condition) composed of 12-s sequences containing 24 visual chunks with written words in different conditions (see Figure 1). For simplicity, we will hereafter refer to visual chunks as single words. As usual in the frequency-tagging paradigm, single words were presented at a constant frequency to keep control of our frequencies of interest (see Frequency spectrum in Figure 1). Specifically, each single word was presented every 0.5 s (0.35 s display + 0.15 s blank screen), resulting in a presentation frequency of 2 Hz. This single-word presentation frequency was shared across sentence and control conditions (see Frequency spectrum in Figure 1). The sentence conditions additionally contained a frequency of interest representing multi-word sentences: as single words were presented every 0.5 s, sentences composed of three words were presented every 1.5 s and thus sentence frequency was at 0.67 Hz (see Frequency spectrum in Figure 1). Each trial started with a fixation cross appearing at the centre of the screen for 0.5 s, immediately followed by stimulus presentation. The inter-trial interval was 6 seconds and participants were invited to blink during this period. Trials were presented in 4 blocks, 2 in Spanish and 2 in Basque, with conditions equally distributed across them. To make sure that participants were paying attention to the stimuli, a recall task was used. At the end of certain trials, written stimuli with a question mark (e.g., ‘hemos vendido una casa?’) appeared on the screen and participants were instructed to report whether or not they had read that exact stimulus during the immediately preceding trial. The presented stimuli varied according to the characteristics of each condition (i.e., NPs were presented in the control conditions, and sentences in the sentence conditions, always in their respective language and real- or pseudo-word version). Answers (yes/no) were provided by pressing two buttons (left-right button position was not counterbalanced across participants). This behavioral task randomly appeared after 33% of the trials and responses could be given during the 5 s post-stimuli presentation. Correct answers were counterbalanced (50 % yes; 50 % no) across trials that required a response.

### Data Acquisition and Pre-processing

MEG data were recorded using a 306-channel Neuromag Vectorview MEG (MEGIN, Helsinki, Finland) located at the BCBL (Donostia-San Sebastián, Spain). Head position was continuously monitored using four Head Position Indicator (HPI) coils. The location of each coil relative to the anatomical fiducials (nasion, and left and right preauricular points) was defined with a 3D digitizer (Fastrak Polhemus, Colchester, VA, USA), in addition to the digitalization of fiducials plus ∼100 additional points evenly distributed over the scalp. MEG recordings were acquired continuously with a sampling rate of 1000 Hz and a bandpass filter at 0.03−330 Hz. Eye movements were monitored with two pairs of electrodes in a bipolar montage placed on the outer canthus of each eye (horizontal electrooculography, EOG), and above and below the right eye (vertical EOG). MEG data were pre-processed off-line using the temporal Signal-Space-Separation (tSSS) method implemented in Maxfilter 2.1 (Elekta-Neuromag) to remove external magnetic noise from the MEG recordings. MEG data were also corrected for head movements and referenced to the initial head position. Bad channels detected during the acquisition were substituted using MaxFilter interpolation algorithms. Subsequent analyses were performed using Matlab R2021B (Mathworks, Natick, MA, USA) and the Fieldtrip toolbox (Oostenveld et al., 2011). First, continuous MEG data were segmented into epochs of 12 s, corresponding to individual trials. Resulting epochs were pre-processed by applying a low-pass filter of 90 Hz. Next, heartbeat and EOG artifacts were detected using Independent Component Analysis (ICA) and linearly subtracted from the MEG data (Infomax algorithm implemented in Fieldtrip). MEG magnetometers (102 sensors) were then discarded, and only planar gradiometers (102 pairs of sensors) were kept for further analysis.

### Data analysis

#### Behavioral task

We were not able to analyze the behavioral responses from the recall task due to technical problems. Specifically, we found a hardware problem in the recording device which was automatically assigning random button-press responses independently of the actual button press across trials. We detected this problem after collecting data from most participants whose responses had extremely short latencies (< 0.005 s). Our behavioral task only occurred in 33% of the trials, was orthogonal to the linguistic manipulations, and was mainly intended to make participants pay attention to the stimuli. Furthermore, as is typical in the frequency-tagging paradigm, we expected neural effects at non-arbitrary frequencies of interest, which in this study clearly differed between the sentence and control conditions. Considering this, we reasoned that the lack of behavioral results should not be critical to interpreting our MEG results.

### Inter-Trial Phase Coherence (ITPC)

To assess the presence of neural synchronization to our frequencies of interest in the MEG data, we first computed ITPC as usual in previous frequency-tagging studies (e.g., (Ding et al., 2017b)). This spectral analysis relies on the frequency-domain decomposition of the whole-trial continuous MEG signal, resulting in a spectrum representing frequency-specific phase consistency strength across trials.

Following previous studies in the auditory domain, we initially removed the first sentence from each trial (i.e., 1.5 s) in the sentence conditions to avoid transient responses evoked by stimulus onset. To make conditions comparable in length, we also removed the first 3 NPs (i.e., 1.5 s) from the control conditions. We then decomposed each trial (10.5 s) into the frequency domain using Fast Fourier Transform (FFT; frequency resolution: 1/10.5 = 0.095 Hz) separately for each planar gradiometer sensor. We extracted the phase from the resulting Fourier coefficients and then computed ITPC following the equation in (Ding et al., 2017b): ITPC(*f*) = (Σ*_k_*cos(θ*_k_*))^2^/*K* + (Σ*_k_*sin(θ*_k_*))^2^/*K*, where *k* is each trial, *K* is the total number of trials, and θ is the phase angle of each complex Fourier coefficient from each frequency ( *f*). Finally, ITPC values from planar gradiometers pairs were linearly combined using Fieldtrip (“sum” method) and the resulting MEG sensors were used for two different ITPC contrasts. First, we averaged ITPC across all MEG sensors to obtain a general measure of phase synchronization at our frequencies of interest in each condition. Second, to assess between-condition ITPC strength differences, we focused on the 12 MEG sensors showing the strongest frequency-specific ITPC values from the grand average between Spanish and Basque sentences separately for the real-word and pseudo-word conditions.

### Phase angle

ITPC reflects phase synchronization strength across trials. However, phase synchronization differences can additionally manifest in terms of distinct temporal dynamics at discrete time points during the trial. This sort of neural phase dynamics cannot be captured by conventional frequency-domain analyses such as ITPC due to their methodological limitations (Zhou et al., 2016; Ding, 2023b; Zhang et al., 2023). Such neural phase dynamics can instead be estimated by assessing the angle orientation of the instantaneous phase derived from time-frequency decomposition of the neural signal. Building on previous reports between phase angle and top-down modulations (Besle et al., 2011; Lakatos et al., 2008; Ten Oever and Sack, 2015; Ten Oever et al., 2024) including abstract structure (Meyer et al., 2016), we examined how the neural phase angle from the sentence frequency (i.e., 0.67 Hz, corresponding to the whole-sentence duration) varied across word boundaries from multi-word sentences. Specifically, we aimed to reveal whether neural activity follows shifted dynamics (approximating anti-phase relationships) between Spanish and Basque sentences with comparable word-level characteristics but reversed abstract structure.

To examine phase angle differences across conditions, we performed time-frequency analysis on the broadband MEG data focusing on the sentence frequency. First, each 12-s trial was further epoched to include its preceding 0.5 s and its subsequent 1.5 s to avoid edge artifacts when computing the frequency-specific time series. MEG data were then low-pass filtered at 30 Hz (filter order = 2) and downsampled to 100 Hz to reduce computation time. Next, separately for each planar gradiometer sensor, we computed the time-frequency representation of each trial using Morlet wavelets only for the sentence frequency (i.e., 0.67-Hz; in steps of 0.01 s) as implemented in Fieldtrip (ft_freqanalysis; cfg.method = ‘wavelet’; cfg.output = ‘fourier’; cfg.width = 3; cfg.gwidth = 3). We then extracted the instantaneous phase of the resulting Fourier coefficients at boundaries corresponding to the exact occurrence of each word (i.e., the offset of the preceding word and the onset of the following word). We discarded the instantaneous phase from the words within the first and last sentences of each trial to avoid trial onset and offset effects, respectively. This analysis was performed for both Spanish and Basque sentence conditions separately between real-word and pseudo-word conditions. To accomplish an unbiased sensor selection, we focused on the planar gradiometer sensors showing the strongest instantaneous phase synchronization strength across the three word boundaries considering the grand average of both languages (separately for real-word and pseudo-word conditions). We thus first implemented the Rayleigh test to evaluate the instantaneous phase synchronization (i.e., non-uniform distribution of phase angles, expressed as z-values) separately for each condition and word boundary using the ‘circ_rtest’ function from the Circular Statistics toolbox (Berens, 2009) in Matlab. Next, we computed the grand average across word boundaries and language conditions. From this grand average, we selected the two sensors with the strongest phase synchronization z-values. We additionally combined the resulting z-values within each planar gradiometer pair using Fieldtrip (“sum” method) only for visualization purposes, as the combination of gradiometers often results in the cancellation of phase angle differences. To assess phase angle differences in the two selected sensors, we computed the circular mean of the phase angles from each participant using the ‘circ_mean’ function from the Circular Statistics toolbox (Berens, 2009) separately for each sentence condition and word boundary.

### Statistical analysis

Statistical significance of linear data (i.e., phase synchronization magnitude) at the group level was examined using distinct analytical methods in Rstudio (Posit Team, 2022; version 2024.4.4.2). Following previous frequency-tagging studies (e.g., (Ding et al., 2016, 2017b), statistical significance of sensor-averaged ITPC was initially tested with a Signal-to-Noise Ratio (SNR) analysis by comparing the ITPC values of each frequency of interest to the mean of the +/- 4 surrounding frequency bins (corresponding to +/- 0.38 Hz; one-sided t-test, uncorrected p-values). Between-condition ITPC differences from strongest MEG sensors were tested using linear mixed-effects models separately for each frequency of interest. Specifically, separately for real-word and pseudo-word conditions, full models included the factor ‘Language’ (Spanish vs. Basque) as fixed effect and ‘Participants’ as random effect. Full models were compared to a reduced model without the ‘Language’ fixed effect using a likelihood ratio test (Chi-Square statistic, Х^2^) and the resulting p-value was used to determine statistically significant differences in the goodness of fit between models. Pairwise comparisons were tested using the *emmeans* function (bonferroni corrected two-sided t-test). Differences in phase synchronization magnitude derived from Rayleigh’s z-values were also tested using comparable linear mixed-effects models.

Statistical significance of circular data (i.e., phase angle distribution) at the group level was examined using two different approaches. First, we implemented the Kuiper test (a circular version of the Kolmogorov-Smirnov test) to assess significant differences between two phase angle distributions in terms of mean location or dispersion using the ‘circ_kuipertest’ function from the Circular Statistics toolbox (Berens, 2009). However, it has recently been suggested that the Watson U^2^ test instead provides more powerful statistical estimates of distribution differences for circular data (Landler et al., 2021). We thus also run the Watson U^2^ test using the ‘watsons_U2_approx_p’ function by Pierre Mégevand (https://github.com/pierremegevand/watsons_u2). We provide both the Kuiper test’s statistic (k) and the Watson’s U^2^ statistic (U^2^) along with their respective p-value (uncorrected).

## Results

### ITPC

The SNR results of sensor-averaged ITPC spectra revealed phase synchronization at the frequency of both single words (2 Hz) and multi-word sentences (0.67 Hz) in all sentence conditions (see Figure 2). For the real-word conditions (Figure 2A), the 2-Hz peak was significant in both Spanish (t(21) = 14.633, p < .001) and Basque (t(21) = 17.434, p < .001) sentences. In addition, the 0.67-Hz peak was also significant in both Spanish (t(21) = 9.2744, p < .001) and Basque (t(21) = 8.885, p < .001) sentences. Similarly for the pseudo-word conditions (Figure 2B), the 2-Hz peak was significant in both Spanish (t(21) = 14.826 p < .001) and Basque (t(21) = 15.634, p < .001) sentences and the 0.67-Hz was also significant in both Spanish (t(21) = 9.4844, p < .001) and Basque (t(21) = 7.8762, p < .001) sentences. All sentence conditions also reflected a significant ITPC peak at 1.33 Hz (p < .001), which corresponds to the harmonic frequency of the 0.67-Hz sentence frequency. Frequency-tagging studies often disregard the interpretation of harmonics as they result from methodological considerations of frequency-domain analysis (see (Ding et al., 2016; Zhou et al., 2016)). In contrast to the sentence conditions, the control conditions only reflected phase synchronization at the frequency of single words (see Figure S1). For the real-word conditions (Figure S1A), the 2-Hz peak was significant in both Spanish (t(21) = 14.992, p < .001) and Basque (t(21) = 20.006, p < .001) but the 0.67-Hz peak was not significant neither in Spanish (t(21) = −1.336, p = .902) nor in Basque (t(21) = −0.427, p = .663). Likewise, for the pseudo-word conditions (Figure S1B), the 2-Hz peak was significant in both Spanish (t(21) = 11.829, p < .001) and Basque (t(21) = 16.205, p < .001) but the 0.67-Hz peak was not significant neither in Spanish (t(21) = 0.413, p = .342) nor in Basque (t(21) = −2.449, p = .988). The SNR results thus confirm the selective presence of phase synchronization to multi-word structures in both Spanish and Basque sentences independently of lexical content (i.e., consistently for both real words and pseudo-words).

**Figure 2.**
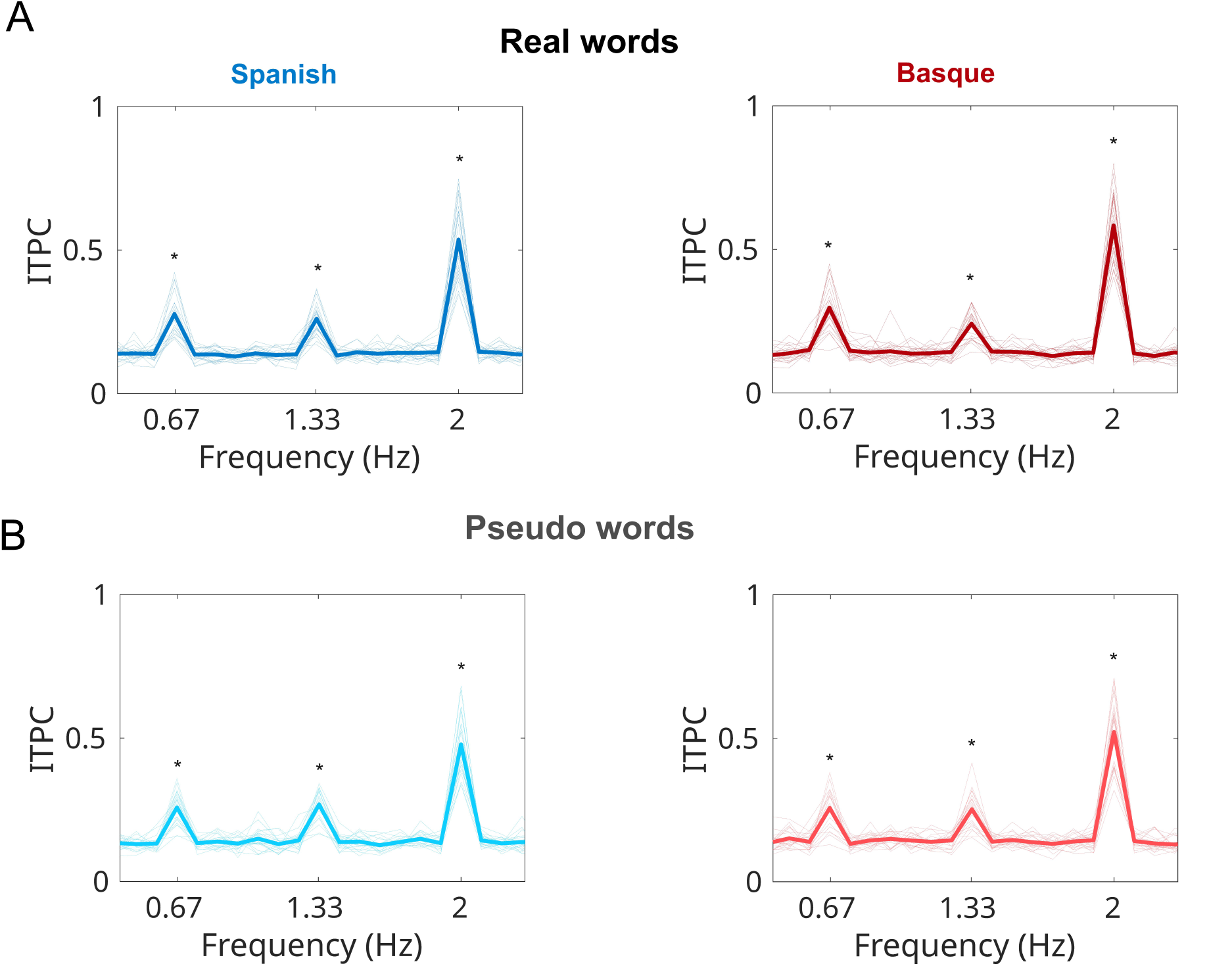
Inter-Trial Phase Coherence (ITPC) spectrum from sentence conditions. ITPC averaged across all sensors, shown separately for each language (Spanish in blue, left; Basque in red, right) and word type (Real words in darker colors at top row; Pseudo words in lighter colors at bottom row). Thinner lines correspond to single-participant average and thicker lines to grand average across participants. X-axis represents frequencies (in Hertz, Hz) and Y-axis ITPC values. Asterisks represent SNR results showing significantly stronger frequency-specific ITPC relative to neighboring (+/-4) frequency bins (see Methods for further details).

The MEG sensors with the strongest ITPC values revealed predominantly similar patterns between Spanish and Basque sentences (see Figure 3). For the real-word conditions (Figure 3A), the 2-Hz ITPC peak corresponding to the single-word frequency was strongest over posterior (occipital and parietal) sensors. This 2-Hz peak was not significantly different between Spanish and Basque conditions (Х^2^(1) = 2.383, p = .123). Moreover, the 0.67-Hz ITPC peak corresponding to the sentence frequency was most prominent in fronto-temporal sensors from the left hemisphere along with posterior (occipital and parietal) sensors relatively leftward lateralized. This 0.67-Hz peak was again not significantly different between Spanish and Basque conditions (Х^2^(1) = 1.273, p = .259). For the pseudo-word conditions (Figure 3B), the sensor-level distribution of both the 2-Hz and 0.67-Hz peaks was highly similar to the real-word conditions. However, while the 2-Hz peak was not significantly different between languages (Х^2^(1) = 1.665, p = .197), we did observe stronger ITPC at 0.67 Hz for Spanish compared to Basque sentences (Х^2^(1) = 7.001, p = .008; t(21) = 2.805, p = .010). To better understand this significant effect, we additionally assessed the contribution of the lexical factor (real words vs. pseudo words) separately for each language. In Spanish sentences, 0.67-Hz ITPC was not significantly different between real-word and pseudo-word conditions (Х^2^(1) = 1.739, p = .187). However, in Basque sentences, 0.67-Hz ITPC was significantly stronger for real words compared to pseudo words (Х^2^(1) = 6.6126, p = .010; t(21) = 2.714, p = .013). Furthermore, in the subset of left-hemisphere fronto-temporal sensors, this ITPC effect (difference for real words minus pseudo words in Basque) was positively correlated Basque proficiency (Spearman rho = 0.43, p = .046) as estimated with BEST scores (de Bruin et al., 2017). These ITPC results thus suggest that phase synchronization strength to Spanish and Basque sentences remains similar with real words, although the presence of pseudo words selectively attenuates phase synchronization strength to Basque (but not Spanish) sentences.

**Figure 3.**
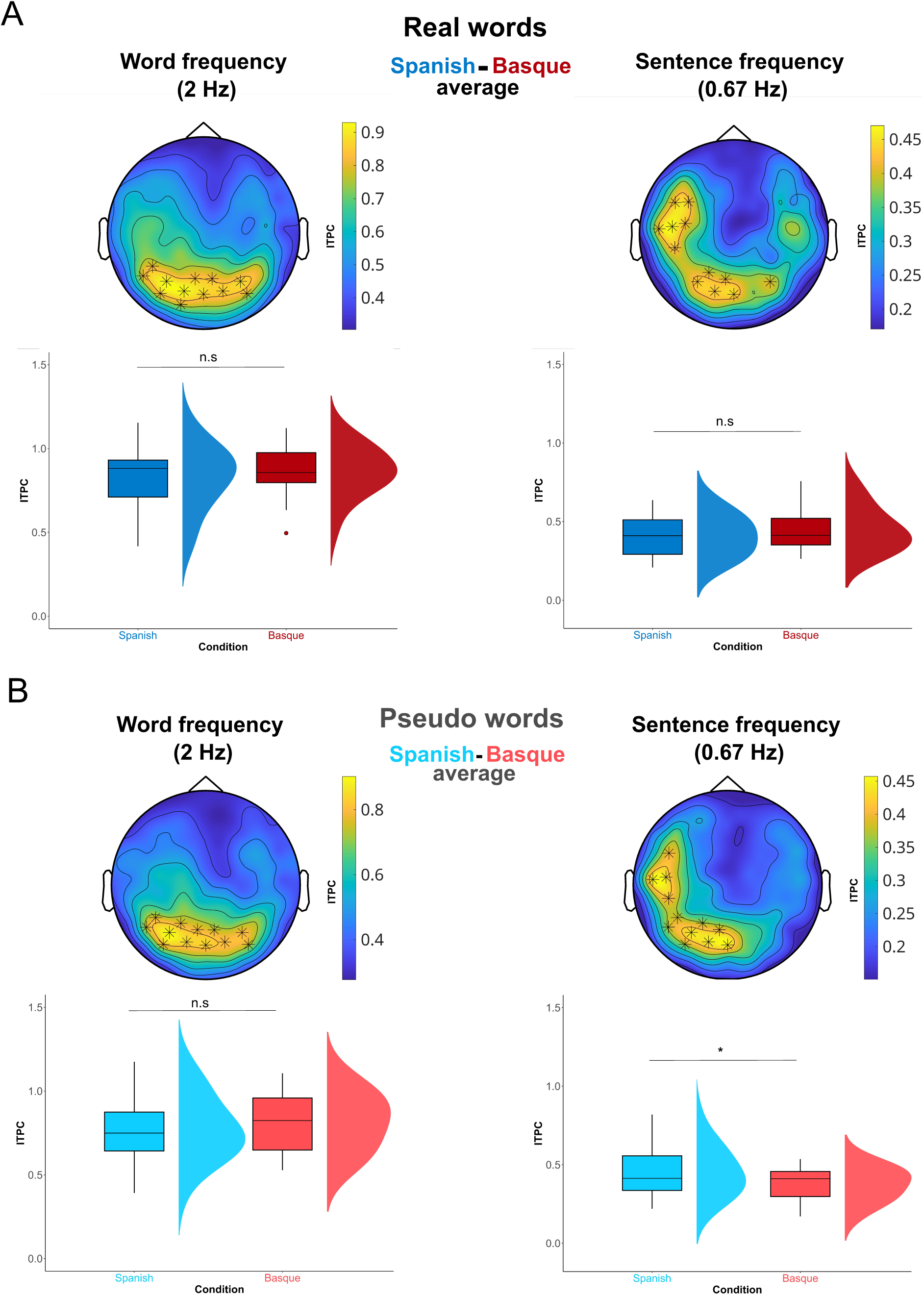
Inter-Trial Phase Coherence (ITPC) results from sentence conditions. Top rows: Sensor-level distribution of ITPC averaged across Spanish and Basque at word frequency (left) and sentence frequency (right) for Real words (A) and Pseudo words (B). Asterisks indicate the 12 sensors showing the strongest ITPC values. Bottom rows: Boxplots displaying the ITPC values averaged across the strongest sensors indicated in corresponding top-row plot separately for Spanish (in blue) and Basque (in red) for Real word (in A, dark colors) and Pseudo word (in B, lighter colors) conditions (central line = median; box outlines = 1st and 3rd quartiles; whiskers = 1.5 interquartile range; raincloud plots display probability density estimates, see Allen et al., 2021). Symbols above boxes represent the results of the statistical analyses testing ITPC differences between Spanish and Basque conditions (asterisk = significant; n.s. = non-significant; see main text for further details).

### Phase Angle

The results of the Spanish-Basque grand average of instantaneous phase synchronization strength across word boundaries (see Figure 4AB) revealed that the strongest sensors were located in a temporal sensor from the left hemisphere (‘MEG0242’ for real-word conditions and ‘MEG1513’ for pseudo-word conditions) and a posterior sensor with a centro-occipital distribution (‘MEG2112’ for both real-word and pseudo-word conditions). Note that this sensor-level distribution is strikingly similar to the ITPC results at the sentence frequency (see Figure 3). In the temporal sensor (Figure 4C), we observed that the circular distribution of phase angles between Spanish and Basque sentences was significantly different across the three word boundaries with real words: boundary 1 (k = 418, p = .001; U^2^ = 0.7798, p < .001), boundary 2 (k = 418, p = .001; U^2^ = 0.7792, p < .001), and boundary 3 (k = 396, p = .001; U^2^ = 0.7792, p < .001); and similarly with pseudo words: boundary 1 (k = 418, p = .001; U^2^ = 0.7720, p < .001), boundary 2 (k = 440, p = .001; U^2^ = 0.7856, p < .001), and boundary 3 (k = 418, p = .001; U^2^ = 0.7474, p < .001). Note that these phase angle shifts cannot be solely due to lexical differences in word class (part-of-speech) between languages at the same word position, as they also emerged at the second word boundary corresponding to a verb in both Spanish and Basque sentences (see Figure 4A). In contrast, in the posterior sensor (Figure 4D), we did not observe significant differences in circular distribution between Spanish and Basque word boundaries with real words: boundary 1 (k = 132, p > .05; U^2^ = 0.0604, p = .6072), boundary 2 (k = 154, p > .05; U^2^ = 0.0604, p = .6072), boundary 3 (k = 132, p > .05; U^2^ = 0.0504, p = .7391); and similarly with pseudo words: boundary 1 (k = 132, p > .05; U^2^ = 0.0726, p = .4772), boundary 2 (k = 132, p > .05; U^2^ = 0.0980 p = .2893), boundary 3 (k = 154, p > .05; U^2^ = 0.0754, p = .4514). Apart from these between-language statistical contrasts, visual comparison in the temporal sensor (Figure 4C) further seems to indicate that phase angle distributions exhibit patterns that are non-identical between word types (i.e., real- vs. pseudo-word sentences) but yet relatively consistent within Spanish and Basque sentences. Phase angle results thus suggest that temporal (but not posterior) sensors reflect shifted temporal dynamics between Spanish and Basque sentences independently of single-word lexical characteristics. To provide a coarse representation of such reversed dynamics, we additionally simulated sinusoidal waves based on the circular mean data at each word boundary in the temporal sensor (see Figure S2). Simulated phase-specific sinusoids show a nearly anti-phase relationship between Spanish and Basque sentences consistently for both real-word and pseudo-word conditions (Figure S2A). Interestingly, both Spanish and Basque sentences displayed highly consistent patterns between word types, with pseudo-words apparently resulting in delayed phase dynamics clearly similar within rather than between languages (Figure S2B). Together, these results provide strong evidence for the presence of shifted phase angle dynamics during the processing of sentences containing reversed abstract structure but nevertheless comparable word-level characteristics.

**Figure 4.**
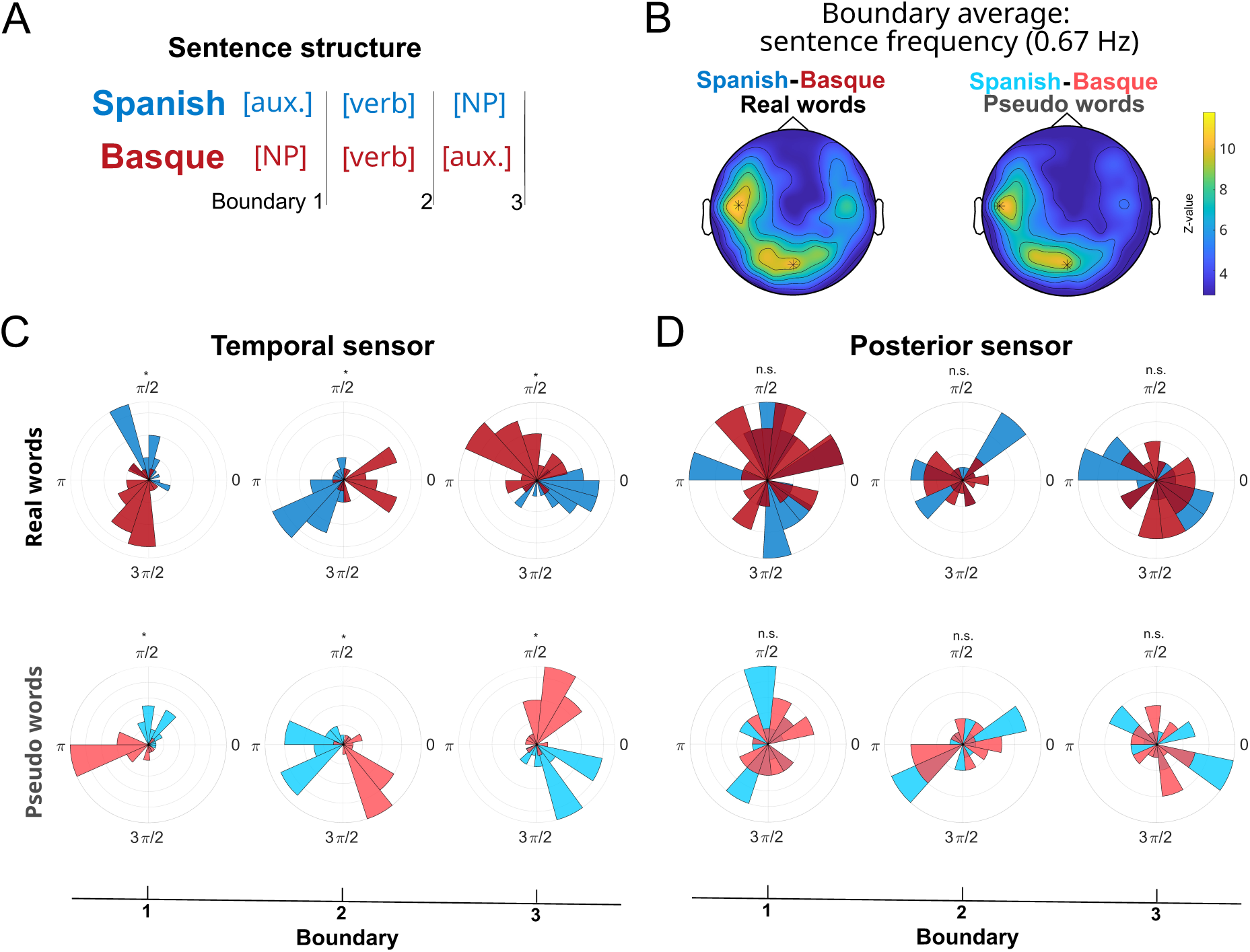
Phase angle results from sentence conditions. A) Schematic representation of extraction of instantaneous phase angle from sentence frequency (0.67 Hz) at three different word boundaries in Spanish (blue) and Basque (red) sentence structures. B) Sensor-level distribution of synchronization strength from sentence-frequency instantaneous phase angle (Z-values from Rayleigh test) averaged across the three word boundaries and both languages separately for Real word (left) and Pseudo word (right) conditions. Asterisks represent the two sensors showing strongest phase synchronization. C) Circular histograms of the circular mean phase angle of each participant (phase values binned into 15 bins, only for visualization purposes) from the two strongest sensors from B): (left-hemisphere) temporal sensor in C) and (central) posterior sensor in D). Top row: Real word conditions (darker colors): Bottom row: Pseudo word conditions (lighter colors), separately for Spanish (in blue) and Basque (in red). Symbols above each histogram represent the results of the statistical analyses testing differences in phase angle distributions between Spanish and Basque conditions (asterisk = significant; n.s. = non-significant; see main text for further details).

A methodological confound that might partially explain the observed phase angle shifts between languages is that both Spanish and Basque sentences contained the constant repetition of identical auxiliary verb words in distinct positions (word 1 in Spanish: *hemos*; word 3 in Basque: *dugu*; see Figure 4A). Stimulus repetition could thus have triggered phase resets at such words, leading to shifted phase dynamics given their different positions between Spanish and Basque sentences. If the reported between-language phase angle shifts were driven by phase resets, it would be reasonable to observe strongest phase synchronization magnitude at such particular word positions in each language. To test this possibility, we assessed statistical differences in instantaneous phase synchronization strength (i.e., Rayleigh’s z-values used to compute the grand average in Figure 4B) across the three word boundaries separately for each condition. For the real-word conditions, we did not observe any significant effect of word position in neither Spanish (Х^2^(2) = 0.1328, p = .936) nor Basque (Х^2^(2) = 2.9271, p = .231) sentences. Similar null effects were observed for the pseudo-word conditions in the Spanish (Х^2^(2) = 5.1942, p = .0745) and Basque (Х^2^(2) = 0.2647, p = .876) sentences. Therefore, the comparable phase synchronization strength across the three word positions is inconsistent with the presence of phase resets at distinct word positions driving the shifted phase angle dynamics between Spanish and Basque sentences.

## Discussion

The current MEG study investigated the relative contribution of structural and lexical components in the neural responses synchronized to the frequency of multi-word structures. To this end, we implemented the frequency-tagging paradigm in the visual modality by presenting Spanish-Basque bilinguals with written words that could be structured into multi-word sentences varying in internal structure (in reversed word orders between Spanish and Basque) and in lexical content (with either real words or pseudo words). We observed robust phase synchronization strength (ITPC) to the frequency of Spanish and Basque sentences containing words with different lexical content, with only Basque exhibiting certain lexical modulations. Notably, we additionally found shifted phase angle dynamics during the processing of Spanish and Basque sentences independently of word-level lexical content. These findings suggest that low-frequency neural synchronization to multi-word structures is dominated by structural rather than lexical components as reflected in two segregated neural dimensions (i.e., frequency and phase). Our MEG results thus extend prior findings on the primary role of structural processing (Ding et al., 2016, 2017a, 2017b; Ding, 2023a, 2025) and advance both theoretical and methodological implications for the neural mechanisms underlying language comprehension.

Low-frequency neural responses to multi-word sentences were mostly driven by structural rather than lexical information. For both Spanish and Basque, we observed robust ITPC at the frequency of multi-word sentences containing either real words or pseudo words. Contrasts across conditions revealed that lexical information only modulated sentence-frequency ITPC for Basque, resulting in attenuated phase synchronization with pseudo words. This cross-linguistic difference cannot solely result from lexical processing alone but rather seems consistent with the language-specific incremental processing dynamics constrained by underlying structure (Levy and Keller, 2013; Husain et al., 2014; Coopmans et al., 2025). Importantly, this ITPC attenuation (real words minus pseudo words in Basque sentences) was positively correlated with Basque proficiency. This suggests that more proficient Basque speakers are better able to use lexical information to support the processing of verb-final sentence structure. Critically, however, the Basque-specific lexical modulation should not be interpreted as evidence for a lexical account of sentence-frequency responses. Instead, it possibly reflects that lexical information can support structural processing when lexical features become available earlier in the input, as in verb-final Basque sentences, where initial arguments containing lexically-informative cues (most prominent with real words than pseudo words) can be used to anticipate sentence structure. The similar sensor-level distribution across words differing in lexical content further supports that sentence-level neural responses are not primarily driven by lexical information. ITPC results are thus generally consistent with a modulatory but not driving role of lexical processing, in line with previous findings (Burroughs et al., 2021; Zhao et al., 2024; Xie et al., 2025). This interpretation is also reinforced by the phase synchronization results derived from the instantaneous phase at each word position. Specifically, phase angle dynamics at the sentence frequency were shifted by structural (Spanish vs. Basque) but not lexical (real words vs. pseudo words) differences consistently across the three words composing sentences. The lower lexical content from pseudo-word sentences only seemed to slightly delay phase temporal dynamics similarly in both languages (as estimated from the simulated signals), which might indirectly index the decreased time-resolved phase synchronization for pseudo-word sentences from previous studies (Kaufeld et al., 2020; Coopmans et al., 2022). Crucially, the phase angle shift was consistently observed between languages even when comparing the identical word class (i.e., the main verb at the second word boundary), suggesting a clear dissociation from word-level lexical processing. The systematic observation of phase synchronization patterns largely independent from word-level information thus provides evidence against lexical accounts (Frank and Yang, 2018; Frank and Christiansen, 2018; Kalenkovich et al., 2022). Instead, our results align with structural accounts regarding the dominant role of structural processing in the neural responses synchronized to multi-word structures (Ding et al., 2016, 2017a, 2017b; Ding, 2023a, 2025) but not necessarily with their assumptions about hierarchical structure (see (Lo et al., 2023)).

Structural information strikingly shifted the phase angle of neural responses between Spanish and Basque sentences containing internally reversed structures. As noted, this sort of anti-phase relationship between languages was found independently of word-level lexical information across lexical content (i.e., real words vs. pseudo words) and word class (i.e., even at the second word position containing the main verb in both languages). Importantly, it was also observed despite the comparable phase synchronization strength across word boundaries, which seems to further indicate a dissociation from phase resets driven by between-language lexical differences at distinct word positions. Together, our findings suggest a selective role of neural phase angle for representing abstract properties internal to reversed structures. This resembles previous top-down modulations of phase angle shifts for segmenting structurally ambiguous sentences into shorter/longer phrases (Meyer et al., 2016; Henke and Meyer, 2021) and extends previous reports on neural sensitivity to other word-order contrasts (Chen et al., 2025) which additionally involved non-structural (e.g., semantic or pragmatic) differences inherent to within-language manipulations. To our knowledge, this is the first demonstration of phase-angle sensitivity to structural features internal to multi-word sentences with identical timescales.

Crucially, our findings have theoretical implications for the characterization of language comprehension mechanisms. Structural accounts generally advocate for frequency-specific mechanisms that exhibit robust phase synchronization to the frequency (i.e., timescales/duration) of multi-word structures (Ding et al., 2016, 2017a, 2017b; Ding, 2023a, 2025). While our results align with such frequency-specific mechanisms considering the largely comparable phase synchronization strength for reversed sentences presented at identical frequencies, our findings also suggest the additional involvement of phase-specific mechanisms that shift the orientation of neural responses depending on the internal properties of multi-word structures. This fundamentally refines how language comprehension mechanisms are currently conceived by segregating distinct structural components into two neural dimensions: frequency coding for the coarser aspects (i.e., timescales/duration) and phase representing the finer-grained aspects (i.e., internal structure) of multi-word structures. This multi-dimensional segregation is compatible with the separation between amplitude and phase hypothesized to instantiate semantic and structural processing, respectively (Martorell et al., 2023). It also provides evidence against recent proposals (Kazanina and Tavano, 2023) suggesting that low-frequency neural responses at the timescales of multi-word structures cannot reliably encode fine-grained aspects of abstract structures. Likewise, our phase angle patterns selectively linked to abstract structure seem to speak in favor of reconsidering previous frequency-tagging evidence about the role of lexical processing based on conventional frequency-domain approaches (Frank and Yang, 2018; Burroughs et al., 2021; Kalenkovich et al., 2022; Lo et al., 2022; Zhao et al., 2024; Chen et al., 2025; Xie et al., 2025). More generally, our findings convincingly demonstrate that core components of language comprehension mechanisms simultaneously manifest across multiple neural dimensions.

The results of the current study also have methodological implications. First, the dissociation between phase synchronization strength and phase angle confirms that conventional frequency-domain measures (e.g., power or ITPC spectra) provide a considerably limited sensitivity to temporal dynamics (for further discussion, see (Zhou et al., 2016; Ding, 2023b; Zhang et al., 2023)). This firmly indicates that crucial insights about language comprehension mechanisms require multi-analytical approaches including conventional frequency-domain measures but also alternative time-frequency measures that consider the temporal dynamics of the neural signal (see also (Martorell et al., 2026)). Second, our study highlights the relevance of cross-linguistic approaches, particularly considering languages differing in typological features (e.g., word order). Our findings thus motivate further cross-linguistic research as a promising approach for broadening the empirical coverage and gaining relevant insights into how language comprehension mechanisms differ in language-specific manners. Third, we provide the first MEG evidence of frequency-tagged responses in the context of structural processing in the visual modality. This replicates a previous EEG study (Pei et al., 2023) which validated the use of written stimuli to investigate structural processing with the frequency-tagging paradigm. In prospective terms, our results strengthen the possibility of using the frequency-tagging paradigm to investigate multi-word language comprehension in populations with whom the use of auditory stimulation is not feasible (e.g., deaf or hearing-impaired populations). Our study also advances potential modality-specific effects in the context of the frequency-tagging paradigm and multi-word language comprehension. Apart from fronto-temporal sensors, posterior sensors located in occipital/parietal regions were also systematically involved in neural synchronization to multi-word sentences. This observation likely reflects the involvement of sensory regions relevant for visual processing, similarly to the findings in the auditory modality showing that acoustic-level and abstract-level effects partially overlap in temporal regions comprising the auditory cortex (Ding et al., 2016; Sheng et al., 2018; Meng et al., 2020). Interestingly, the phase angle effect was only observed in fronto-temporal but not posterior sensors. This suggests that, at least in the visual modality, regions reflecting neural synchronization to abstract structure might underlie considerably different dynamics. Further studies are required to shed light on how auditory and visual modality differentially impact the neural synchronization to multi-word structures.

Our study also has certain limitations. First, we were not able to analyze the results of the behavioral task due to technical problems. Although these behavioral data could have allowed us to potentially discard participants paying low attention to the stimuli, we emphasize that this task only occurred on 33% of the trials and was orthogonal to the linguistic manipulations. Therefore, in this case, we think that the lack of behavioral data may not compromise the here reported MEG results. Second, our participant sample was relatively small (22 participants included in the reported analyses) due to resource constraints (Lakens, 2022). Although our relatively small sample size makes our findings hypothesis-generating rather than conclusive, this should not undermine the reliability of our results. More concretely, most frequency-tagging studies have systematically replicated effects at the frequency of multi-word structures with fewer participants using MEG (e.g., 8 in (Ding et al., 2016); 18 in (Sheng et al., 2018); 19 in (Meng et al., 2020); 20 in (Chen et al., 2025)) or even using EEG (e.g., 16 in (Ding et al., 2017b); 20 in (Burroughs et al., 2021)) which has a lower signal-to-noise ratio for superficial cortical sources (e.g., (Goldenholz et al., 2009)). Finally, our MEG results were only based at the sensor level. Although gradiometers provide a good mapping of superficial cortical sources (Hämäläinen et al., 1993), this makes it impossible to make strong claims about the involvement of specific brain regions. Source-localized MEG signals could have instead revealed more subtle spatial dissociations across fronto-temporal regions, as found in previous frequency-tagging studies between different structures (e.g., (Sheng et al., 2018)). These limitations should be resolved in future studies.

## Conclusion

Our MEG results provide converging cross-linguistic evidence that low-frequency neural responses largely synchronize to structural rather than lexical features during sentence comprehension. We observed robust phase synchronization strength to the frequency of internally reversed structures between Spanish (head-initial) and Basque (head-final) sentences, with certain lexical modulations (real words > pseudo words) depending on language-specific structural features. Crucially, time-resolved analyses revealed shifted phase angle dynamics between the internally reversed structures of Spanish and Basque sentences independently of word-level lexical characteristics. Together, these findings suggest that neural synchronization to multi-word structures is largely driven by distinct structural features operating via two segregated neural dimensions: frequency coding for the coarser aspects (i.e., timescale/duration) and phase representing the finer-grained aspects (i.e., internal structure) of multi-word structures. Our MEG findings thus provide key insights into the core components of the neural mechanisms underlying language comprehension.

## Data and code availability

The pre-processed data and code supporting the findings of this study will be made accessible after publication

## Acknowledgments

We thank BCBL lab staff for help with participant recruitment, Manex Lete for assistance in data collection, and Craig Richter for support in experiment setup.

## Author contributions

conceptualization: J.M., S.M., N.M.,

methodology: J.M., N.M.

formal analysis: J.M.

writing – original draft: J.M., N.M.

writing – review & editing: J.M., S.M., P.M. P-A., M.C., N.M.

investigation, data curation, visualization: J.M.

funding acquisition: S.M., P.M. P-A., M.C., N.M.,

supervision: N.M.

## Supplementary materials

**Table S1.** Descriptive statistics of the psycholinguistic variables of the experimental stimuli.

| Language | Word class | Variable | Mean (SD) | Min | Max |
| --- | --- | --- | --- | --- | --- |
| Real words |  |  |  |  |  |
| Spanish | NP | Frequency | 0.08 (0.1) | 0.002 | 0.65 |
|  |  | Length | 9.1 (1.3) | 5 | 11 |
|  | Verb | Frequency | 0.016<br>(0.023) | 0.004 | 0.16 |
|  |  | Length | 8.3 (1.6) | 4 | 12 |
| Basque | NP | Frequency | 0.08 (0.11) | 0.002 | 0.65 |
|  |  | Length | 7.05 (1.95) | 3 | 12 |
|  | Verb | Frequency | 0.21 (0.35) | 0.01 | 0.35 |
|  |  | Length | 6.59 (1.64) | 2 | 12 |
| Pseudo words |  |  |  |  |  |
| Spanish | NP | Length | 9.1 (1.3) | 5 | 11 |
|  | Verb | Length | 8.4 (1.5) | 5 | 12 |
| Basque | NP | Length | 7.05 (1.95) | 3 | 12 |
|  | Verb | Length | 6.59 (1.64) | 2 | 12 |

*Note. Frequency values are expressed per thousand of words only for real words. Length is measured in number of characters. NP = Noun Phrase; SD = standard deviation; Min = minimum; Max = Maximum.*

**Figure S1.**
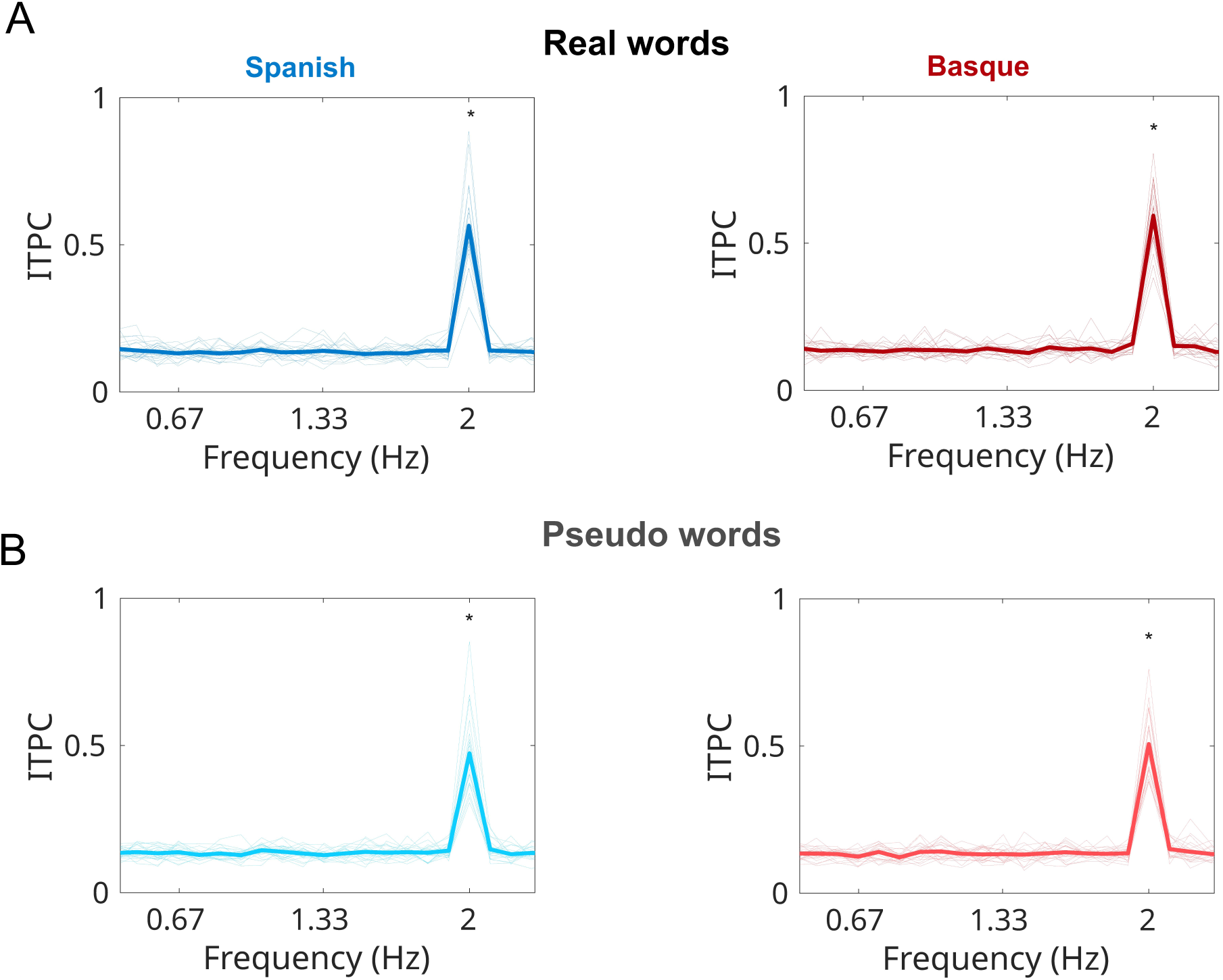
Inter-Trial Phase Coherence (ITPC) spectrum from control conditions. ITPC averaged across all sensors, shown separately for each language (Spanish in blue, left; Basque in red, right) and word type (Real words in darker colors at top row; Pseudo words in lighter colors at bottom row). Thinner lines correspond to single-participant average and thicker lines to grand average across participants. X-axis represents frequencies (in Hertz, Hz) and Y-axis ITPC values. Asterisks represent SNR results showing significantly stronger frequency-specific ITPC relative to neighboring (+/-4) frequency bins (see Methods for further details).

**Figure S2.**
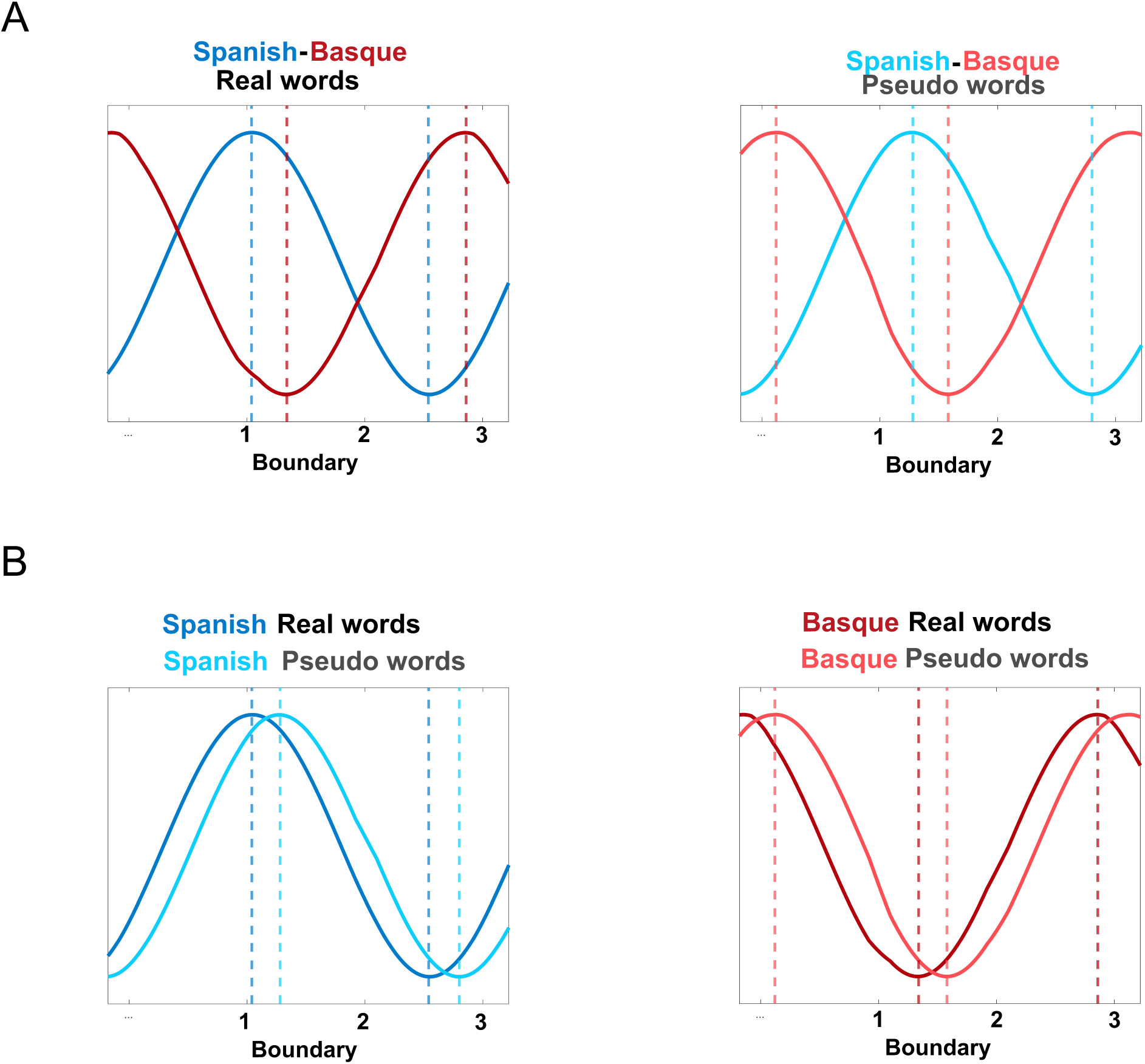
Simulations of phase angle dynamics. **A)** Simulated sinusoidal waves based on phase angles values (circular mean across participants) from each word boundary in temporal sensor (see Figure 4C) for Spanish (in blue) and Basque (in red) and for Real words (left, in darker colors) and Pseudo words (right, lighter colors). **B)** same data as in A with within-language contrasts between Real and Pseudo word conditions for Spanish (left) and Basque (right) conditions. Vertical dashed lines indicate the peaks and troughs of each simulated wave (see main text for further details).

